# Novel structure and composition of the unusually large germline determinant of the wasp Nasonia vitripennis

**DOI:** 10.1101/2024.11.01.621563

**Authors:** Allie Kemph, Kabita Kharel, Samuel J. Tindell, Alexey L. Arkov, Jeremy A. Lynch

## Abstract

Specialized, maternally derived ribonucleoprotein (RNP) granules play an important role in specifying the primordial germ cells in many animal species. Typically, these germ granules are small (∼100 nm to a few microns in diameter) and numerous; in contrast, a single, extremely large granule called the oosome plays the role of germline determinant in the wasp *Nasonia vitripennis.* The organizational basis underlying the form and function of this unusually large membraneless RNP granule remains an open question. Here we use a combination of super-resolution and transmission electron microscopy to investigate the composition and morphology of the oosome. We show that the oosome has properties of a viscous liquid or elastic solid. The most prominent feature of the oosome is a branching mesh-like network of high abundance mRNAs that pervades the entire structure. Homologs of the core polar granule proteins Vasa and Oskar do not appear to nucleate this network, but rather are distributed adjacently as separate puncta. Low abundance RNAs appear to cluster in puncta that similarly do not overlap with the protein puncta. Several membrane-bound organelles, including lipid droplets and rough ER-like vesicles, are incorporated within the oosome, whereas mitochondria are nearly entirely excluded. Our findings show that the remarkably large size of the oosome is reflected in a complex sub-granular organization and suggest that the oosome is a powerful model for probing interactions between membraneless and membrane-bound organelles, structural features that contribute to granule size, and the evolution of germ plasm in insects.

## Background

Primordial germ cells are generally specified early in development to prevent somatic differentiation and ensure the successful production of gametes. In many sexually reproducing organisms, germ cells are among the first to be specified via their uptake of a specialized cytoplasm consisting of maternally deposited mRNAs and proteins, termed ‘germ plasm’ [1]. The germ plasm is typically characterized by numerous membraneless complexes that are highly concentrated in mRNA and RNA-binding proteins called germ granules. While there is significant variability in the contents of these granules [2, 3], there are some highly conserved factors including mRNAs encoding *nanos* homologs, and conserved core proteins of the Vasa-Ddx4, Tudor, and Argonaute-Piwi families [4]. The germ plasm nucleator varies across species, but these proteins often share intrinsically disordered regions (IDRs) believed to play an important role in driving liquid-liquid phase separation [5-7].

Among holometabolous insects, germ plasm nucleation is dependent upon the gene *oskar* and its loss correlates with shifts to the alternative zygotic induction specification strategy [8]. *osk* was originally only detected in the dipteran lineage where it has been studied extensively in the model organism *Drosophila melanogaster* [9, 10]. An *osk* ortholog was eventually discovered in the wasp *Nasonia vitripennis*, where it was shown to have a conserved (if highly modified) role in germ plasm nucleation [8]. More recently it was found that *osk* arose early in insect evolution and originally played no role in germline determination [11-13]. In both *Drosophila* and *Nasonia, osk* mRNA localizes to the posterior pole during oogenesis, where it is translated and functions upstream of the other core assembly proteins Vasa and Tudor [8, 14]. In *Drosophila*, these proteins form many small 500nm ribonucleoprotein complexes called polar granules, phase-transitioned particles with both liquid and hydrogel-like properties [15]. These granules bind, protect, and regulate the expression of germline mRNAs. With the advent of super-resolution imaging techniques, it has emerged that polar granule-localized mRNAs accumulate in distinct homotypic clusters in the context of these protein-based granule [16, 17].

In stark contrast, the germ plasm of *Nasonia* takes the form of a single large particle, forty-fold larger than the individual polar granules. How this unusually large size for a membraneless intracellular structure is achieved and maintained is of very high interest. We have previously shown that the mRNA profile of the oosome is quite different from the polar granules, only sharing a handful of mRNA molecules [3]. Our recent examination of the protein composition and arrangement of the oosome revealed granules of Osk/Vasa/Aubergine that are similar in size to polar granules distributed throughout the oosome [18]. However, they showed limited colocalization suggesting that they do not form the same type of structures as the polar granules. In addition we found that Tudor protein forms a novel shell-like distribution around the surface of the oosome, which has no counterpart in the fly [18].

Several unanswered questions about the structure and composition of the oosome still remain, that we begin to address here. First, we explore the development of its shape throughout the pre-blastoderm stage to gain insights into its behavior and biophysical properties. Additionally, since the nucleation of homotypic clusters of mRNAs is a major, functionally important feature of polar granules, we explore how mRNAs of different enrichment levels are distributed in the oosome relative to each other, and to the core protein components. Finally, due to the pronounced enrichment of mitochondria in *Drosophila* germ plasm [19], and the demonstrated importance of membrane-bound organelles in regulating polar granule morphology [20], we’ve also examined the distribution of subcellular components in and around the oosome. Our analyses identified a clear pattern in oosome morphology, unexpected organelle interactions, and revealed a novel arrangement of germ plasm mRNAs.

## Results

### Dynamic oosome morphology during early embryogenesis

We previously observed that the oosome underwent dramatic changes and migration in the early embryo prior to cellularization [8]. We examined oosome behavior during this period which encompasses the first 3 hours of development, comprising the first seven syncytial nuclear divisions and most of the pre-blastoderm stage prior to pole cell budding, to gain insights into the material properties of the oosome and the forces acting upon it. For the first hour of development, the oosome is attached to the posterior cortex (Figure 1A), where it was assembled during oogenesis. After the maturation of the female pro-nucleus and the first syncytial nuclear division, the oosome detaches and migrates anteriorly as far as 50% egg length through nuclear cycles 2 and 3. The oosome then moves posteriorly in cycles 4 and 5, until it reassociates with the cortex during the sixth syncytial nuclear division. It finally becomes flattened against the posterior cortex during the seventh cycle, just before naïve nuclei arrive and pole cell budding initiates (Figure 1A).

**Figure 1:**
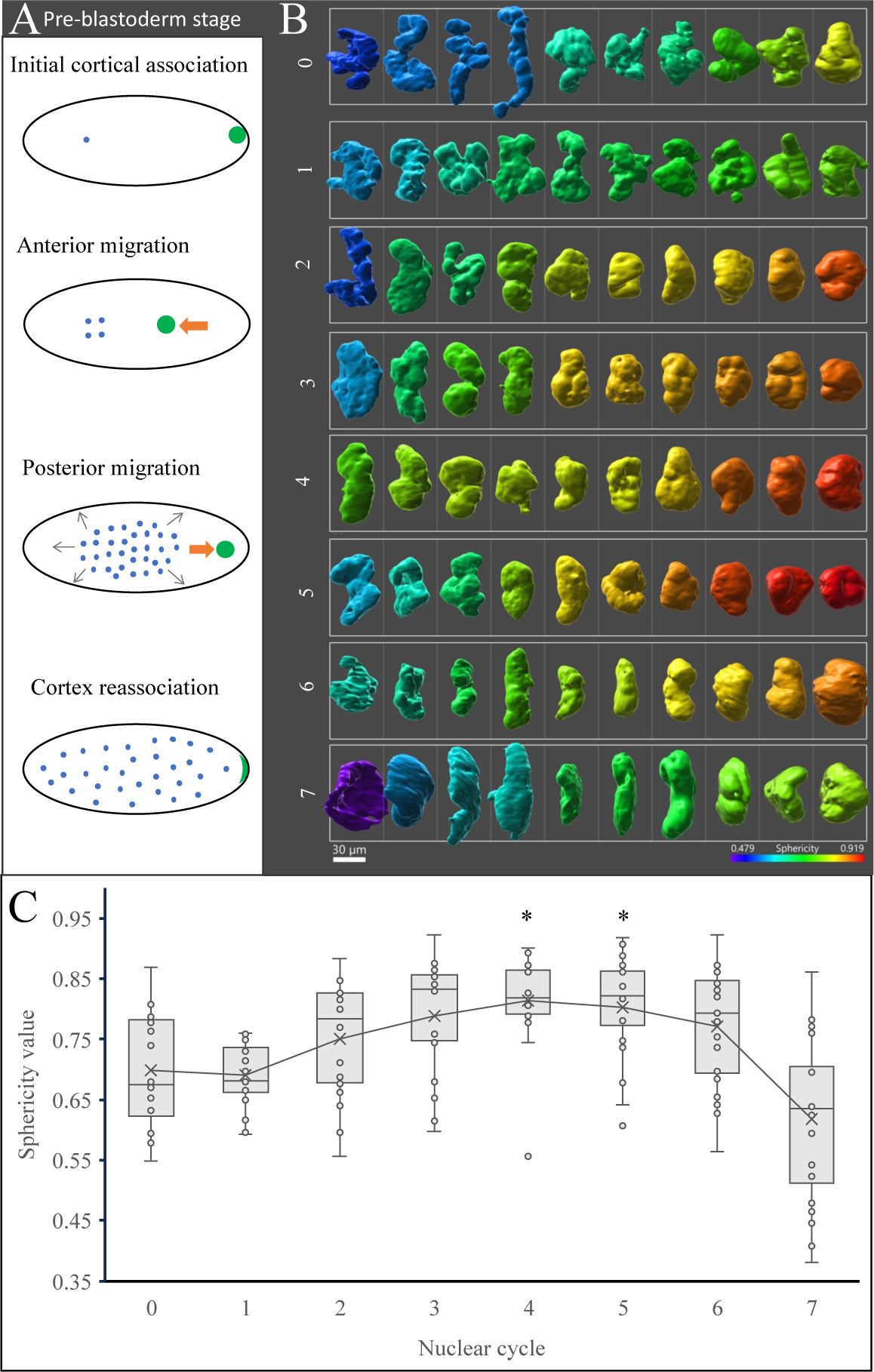
Oosome morphology is dynamic with lower sphericity at cortical association than throughout migration. (A) A diagram of oosome behavior during the pre-blastoderm stage. The oosome is shown in green and nuclei in blue. The orange arrow indicates the direction of oosome migration, and the grey indicates nuclei movement. (B) A gallery plot of oosome sphericity across syncytial nuclear cycles. Color indicates relative sphericity. (C) A boxplot of sphericity values across syncytial nuclear divisions. n=20 for each nuclear cycle. Asterisks indicate significance relative to nuclear cycle 0, as assessed by Dunn’s Test.

Oosomes from each nuclear division cycle were imaged with spinning-disc confocal microscopy at high z-resolution using fluorescent probes against the abundant and ubiquitous *Nv-osk* transcript to delineate the borders of the oosome. Surfaces were generated around the reconstructed oosome projections and were measured for sphericity. We suspect that the combination of the forces driving the oosome’s migration, together with resistance to this force by remaining cortical attachments are responsible for the deformation of the oosome into complex irregular shapes with low sphericity observed in NC0 as the oosome begins detaching from the posterior cortex (Figure 1B & C). Once the oosome is completely detached from the cortex, it exhibits gradually increasing sphericity as it is suspended in the central column of the embryonic cytoplasm. This behavior is indicative of relaxation to minimize surface tension, which is a property of either a viscous liquid or an elastic solid [21, 22]. Finally, as the oosome is pushed back against the posterior cortex it flattens and spreads into a lens-like shape (Figure 1B & C), resembling membrane wetting behavior of a liquid droplet [23] or the deformation of an elastic solid. Taken together, the dynamic morphology of the oosome throughout the pre-blastoderm stage suggest that it displays properties of either a viscous liquid or an elastic solid. The development of fluorescently tagged proteins and associated methods like FRAP in *Nasonia* will be required to make a final determination of the physical properties of the oosome.

### Distribution of mRNAs within the oosome

*Drosophila* germ plasm contains polar granules which include at least two distinct ribonucleoprotein complexes: germ granules and founder granules [17, 24], The latter are responsible for the decay of germ plasm-localized *osk* mRNA, which is toxic for pole cell development [24]. The germ granules are the major functional component of the *Drosophila* germ plasm. They consist of a conserved core set of proteins (Osk, Vas, Tud, and Aub) that are distributed throughout each granule in complex patterns of more or less colocalization [16, 25, 26]. The protein core is decorated by homotypic clusters of mRNAs which occupy specific positions within the granule that are correlated with their expression levels [16, 17, 27].

The core protein components of the germ granules are conserved as constituents of the oosome in *Nasonia*. Our recent super-resolution imaging of the pre-blastoderm oosome revealed that Nv-Osk, Nv-Vas, and Nv-Aub accumulate in only partially co-localized puncta that are distributed throughout the oosome (Fig 2B’’ & C’’) [18]. Nv-Tud differs in that it is highly concentrated around the oosome periphery, with some puncta dispersed interiorly (Figure 2A’’) [18]. Whether this distinct protein arrangement impacts the distribution of the dozens of transcripts that are localized to the oosome remains unknown [3].

**Figure 2:**
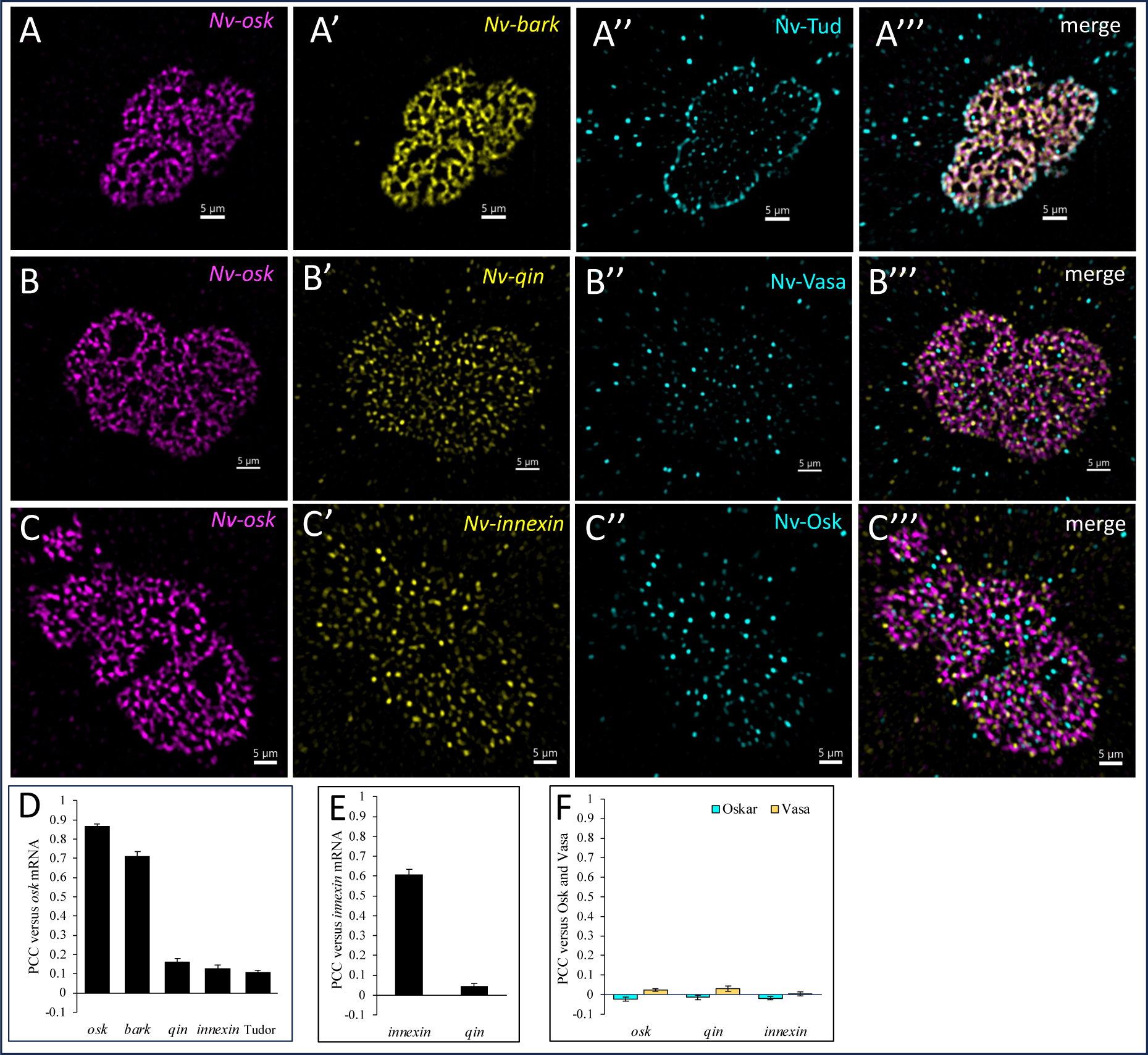
Oosome RNAs exhibit distinct distribution patterns depending on expression levels in the oosome, and show little co-localization with the core germ plasm proteins. (A-A’’’) Single plane confocal SRRF image showing the expression of Nv-osk (magenta), Nv-bark (yellow) and Nv-Tud (cyan) in oosomes. (B-B’’’) Single plane confocal SRRF image showing the expression of Nv-osk (magenta), Nv-qin (yellow) and Nv-Vasa (cyan) in oosomes. (C-C’’’) Single plane confocal SRRF image showing the expression of Nv-osk (magenta), Nv-innexin (yellow) and Nv-Osk (cyan) in oosomes. (D-F) PCC co-localization coefficients detected between oosome mRNAs stained with stellaris probes and antibody-labelled proteins (mean +/– standard error of mean, n=6-12). All scale bars, 5 μm.

To address this gap, we selected four oosome-localized transcripts – *Nv-oskar (Nv-osk)*, *Nv-barkbeetle (Nv-bark)*, *Nv-innexin*, and *Nv-qin* for super-resolution fluorescent *in situ* hybridization. *Nv-osk* and *Nv-bark* are two of the most highly enriched transcripts in the oosome and are expressed at levels an order of magnitude higher than typical oosome genes, represented by *Nv-qin* and *Nv-innexin* [3]. The high abundance transcripts *Nv-bark* and *Nv-osk* are distributed in apparently overlapping mesh-like networks that pervade the oosome (Figure 2A, 2A’, & 2A’’’). There appear to be regions of relatively high, and relatively low intensity along those networks, indicating a variable intensity in clustering of these mRNAs. Such pattern has not been reported to our knowledge in other germ granules where homotypic clustering of mRNAs has been observed. In contrast, the lower abundance transcripts (*Nv-qin* and *Nv-innexin)* cluster into distinct puncta that are distributed relatively diffusely within the oosome and appear to be juxtaposed with the *Nv-osk/bark* network (Figure 2B, B’, C, & C’).

To quantify our qualitative observation about the relative positions of the four transcripts with respect to one another, we performed colocalization analyses of each transcript relative to the *Nv-osk* network. To determine the upper limit of co-localization for our analysis we labeled *Nv-osk* with a mix of stellaris CALFluor®Red 590-labelled probes and Quasar®670-labelled probes and measured a PCC of 0.87± 0.03 (Figure 2D). *Nv-osk* and *Nv-bark* highly co-localize within the oosome with a PCC of 0.71 ± 0.05, where they appear to co-label a mesh-like network running throughout the oosome interior (Figure 2A’’’ & D). The punctate *Nv-innexin* and *Nv-qin* only weakly co-localize with *Nv-osk* with a PCC of 0.16 ± 0.05 and 0.13 ± 0.05, for *Nv-qin* and *Nv-innexin* respectively (Figure 2B-D). *Nv-innexin* and *Nv-qin* also show no co-localization with each other with a PCC of 0.05± 0.0, supporting the presence of homotypic RNA clustering in the oosome (Figure 2E).

We next investigated how these transcripts interact with the protein components of the oosome. As previously described, Nv-Osk and Nv-Vasa, are distributed as relatively diffuse puncta throughout the oosome, while Nv-Tud is highly concentrated at the oosome surface (Figure 2A’’, B’’, & C’’) [18]. Neither Nv-Osk (PCC of -0.02± 0.03), nor Nv-Vasa (PCC of 0.02± 0.03) showed any co-localization with the *Nv-osk* mRNA network (Figure 2B, C, & F). Nv-Tud showed only weak co-localization (PCC of 0.11 ± 0.04), which may reflect a greater likelihood of the continuous network of Nv-Tud to intersect or overlap with the extensive network of *Nv-osk* (Figure 2A’’’&D). Surprisingly, we found that the punctate mRNAs *Nv-innexin* and *Nv-qin* also showed no co-localization with these internal protein granules (Figure 2F). Both *Nv-innexin* and *Nv-qin* are maintained in *Nasonia* pole cells and their lack of co-localization suggests these internal Nv-Osk/Vasa granules are not as essential as germ granules in transcript germ plasm localization and transport to pole cells [3].

Non-homogenous RNA distribution is a common feature in germ granules and our results thus far suggest that is also the case in *Nasonia*. The arrangement of RNA into homotypic clusters has been described in germ granules of several model organisms (*Drosophila*, *C. elegans* and zebrafish) [2, 16, 17, 28, 29], and the punctate organization and lack of co-localization of *Nv-innexin* and *Nv-qin* support the existence of these clusters within the oosome as well (Figure 2B’, C’, & E). Although our co-localization results suggest that *Nv-osk* and *Nv-bark* appear to be distributed along the same network (Figure 2D), they may still form distinct clusters that cannot be resolved with our current optical capabilities. Indirect evidence for this is our previous observations that *Nv-osk* mRNA is rapidly degraded in the pole cells [8], while *Nv-bark* remains at high levels in the primordial germ cells throughout embryogenesis [3]. We have observed that this difference in regulation is apparent much earlier in the oosome, with *Nv-osk* signal showing significant reduction in the intact oosome prior to pole cell formation (Figure S1).

### Fragmentation and stability of the oosome

In most species, germ plasm consists of many small amorphous ribonucleoprotein particles that are of a relatively uniform size (often less than a few micrometers) [30, 31], but granule size can increase both naturally over time through fusion events [31]. as well as artificially through protein knockdown [20, 32]. For example the MEG-3 protein clusters that form a shell around P granules in *C. elegans* are hypothesized to act as a Pickering agent, which is supported by the finding that MEG-3 knockdown results in fewer, larger granules [32]. The oosome somehow naturally bypasses the size constraints generally observed in germ granules forming a uniquely large germ plasm complex. Previous observations showed that the disruption of Tudor, which forms a shell around the oosome (Figure 2A’’, [18]) results in smaller oosomes [8], suggesting that Tudor may stabilize the large size of the oosome, which contrasts with the role of the shell surrounding P granules.

In addition to the singular large oosome, we found smaller aggregates of germ plasm material at the posterior pole, which we named satellite particles. The composition of these satellite particles appears to be identical to the oosome. The network of *Nv-osk/bark* mRNA punctuated with scattered spots of lower abundance RNAs and core proteins (e.g., Nv-Osk, Nv-Vasa) also appears to be maintained (Figure 3A & B). Investigating the origin and fate of these satellite particles and the oosome should provide insights into the mechanisms governing germ granule size.

**Figure 3:**
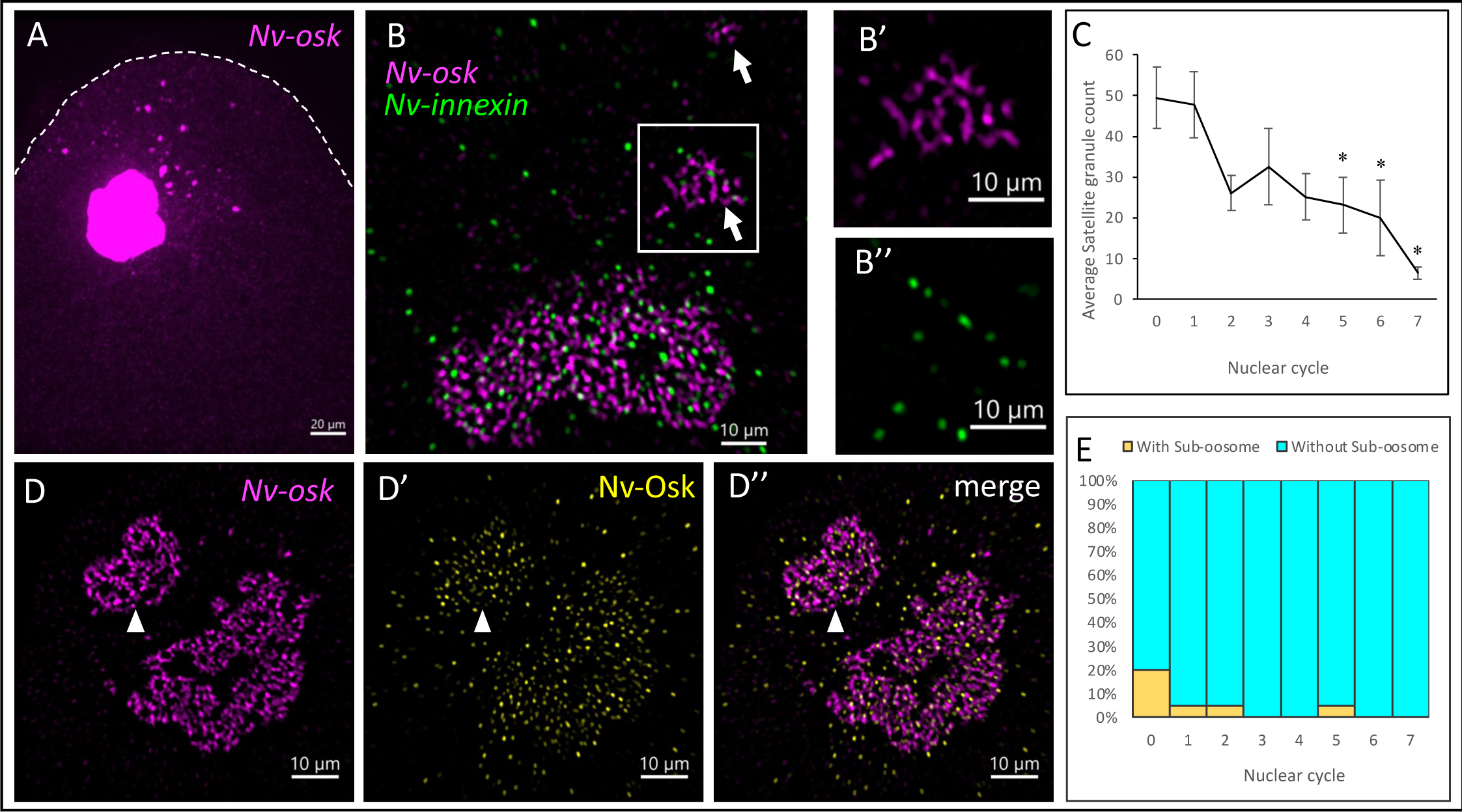
Smaller germ plasm aggregates (satellite particles) exist at the posterior cortex orbiting the oosome and decrease throughout the pre-blastoderm stage. **(A)** Single plane confocal image of the *Nv-osk* (magenta) labeled germ plasm particles in an early-stage embryo posterior. **(B)** Single plane confocal SRRF image of the *Nv-osk* (magenta) and *Nv-innexin* (green) labeled germ plasm particles. White arrows indicate satellite particles **(B’-B’’)** Enlargement of the satellite particle indicated by the white box in B. **(C)** Quantification of the average number of satellite particles across syncytial nuclear cycles. Values shown are mean ± S.E.M; n = 20 embryos per nuclear cycle. Asterisks indicate statistical significance relative to before the first syncytial nuclear cleavage, as assessed by Dunn’s Test. **(D-D”)** Single plane confocal SRRF image of a sub-oosome labeled with *Nv-osk* (magenta) and Nv-Osk (yellow). White arrowhead indicates sub-oosome. **(E)** Quantification of sub-oosome frequency across syncytial nuclear cycles. For each nuclear cycle n=20 embryos. Scale bar in A is 20 μm. All other scale bars are 10 μm.

Satellite particles are variable in size, ranging from similar in size to polar granules to others nearly as large as the oosome, with a continuum of intermediate sizes between those extremes. The number of satellite particles was variable across embryos, however, within that variability we could detect a clear trend of decreasing satellite particle count over developmental time (Figure 3C). The decreasing number of satellite particles suggests they are either fusing together and with the main oosome over time, or that the smaller particles are dissolving in the bulk cytoplasm. We cannot currently distinguish between these possibilities.

Another open question is the origin of the satellite particles. Their significantly higher numbers in the earliest division cycles suggests that they either assemble concurrently with the oosome during oogenesis, or that they arise through fragmentation of the oosome under the strong deforming forces that are acting upon it at this stage (Figure 1). In the case of the larger particles, a subset we have termed sub-oosomes (volume 20% or greater than the main oosome), we propose that they are primarily the result of fragments breaking free from the main oosome, and that these in large part re-fuse with the oosome in later stages. In support of this proposal, sub-oosomes are found with the highest frequency in the earliest nuclear cycles and most closely resemble oosome architecture and have even been observed to share the distinct Tudor-shell distribution (Figures 3D, E & S2A’). Further, we observe putative intermediate cases in the earliest nuclear cycles where a large fragment of the oosome is only tenuously connected to the rest and appears to be in the process of detaching to form a sub-oosome (Movie S2).

### Ultrastructural analysis of the oosome

Our fluorescence based analyses have suggested that the underlying structure of the oosome may be more heavily RNA based than *Drosophila* germ granules [20], and perhaps germ granules from other species. To gain a higher resolution understanding of the structure of the oosome, we examined it with transmission electron microscopy (TEM). The main substance of the oosome appears to be an electron-dense fibro-granular network with pockets of lower electron density material distributed throughout (Figure 4A-A’). This arrangement is strikingly reminiscent of the mesh-like organization of the highly enriched mRNAs *Nv-bark* and *Nv-osk* surrounding areas of low to no signal (Figures 2A’’’ & 4A’). This further suggests that the underlying architecture of the oosome is composed primarily of a mesh-like network of RNA.

**Figure 4:**
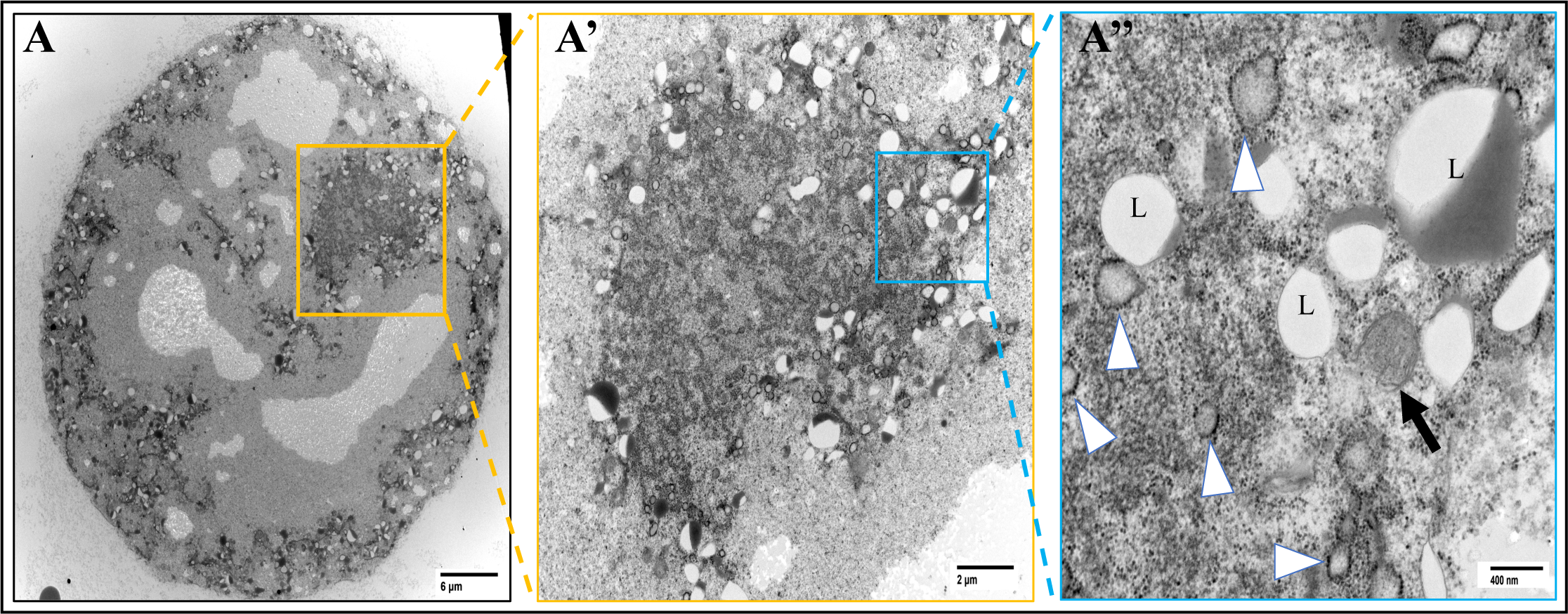
Oosome ultrastructure. **(A)** Electron micrograph of an embryo posterior cortex containing the oosome (yellow box surrounding). Scale bar 6 μm. **(A’)** Enlargement of the yellow box containing the oosome in A. Scale bar 2 μm. **(A’’)** Enlargement of a peripheral portion of the oosome indicated by the blue box in A’. Lipid droplets indicated with L, mitochondria with a black arrow, and RAVs with white arrowheads. Scale bar 400nm.

Germ granules are associated with membrane-bound organelles in multiple species [33, 34], and TEM revealed that the oosome conforms to this broad pattern. Most immediately evident are the putative lipid droplets concentrated to the periphery of the oosome as well as frequently within it (Figure 4A”). There are no identifiable Golgi-like structures, nor are there canonical tubular or sheet-like endoplasmic reticulum structures. However, we did observe spherical membrane-bound organelles studded with what appear to be ribosomes that were distributed both within the oosome and throughout the bulk cytoplasm. These are highly reminiscent of the newly defined ER-like organelle compartment called Ribosome-Associate Vesicles (RAVs) (Figure 4A” white arrowheads) [35]. These were recently described in secretory cell lines and have retroactively been identified in various organisms and across different cell types [35]. Although they have not previously been observed in embryonic germ plasm, similar structures (termed RER vesicles) were described in the “pseudo-pole plasm” in the ovary of an earwig (Dermaptera) [36, 37], These ribosome-associated vesicles have been proposed to transport active ribosomes to distal neurites to promote local translation [38]. Although it is unclear if this role in promoting local translation extends to the context of the large, dense structure of the oosome.

Surprisingly, we found very little association between the oosome and mitochondria. Mitochondria are occasionally seen adjacent to the oosome, and almost never within it. They do not appear to be enriched in any way around the oosome. This differs from *Drosophila melanogaster,* where mitochondria are strongly enriched around polar granules, and are often in direct contact. This may suggest alternative energy requirements or sources between the oosome and polar granules. To confirm and expand on these observations we next visualized both structures within and around the oosome using fluorescence microscopy.

### Interactions of the oosome with membrane-bound organelles

Although RAVs have not been found in embryonic germ plasm of any other animal species, *Drosophila* polar granules have been found to contact conventional endoplasmic reticulum [20, 39]. To further validate that the unusual vesicles we see in the *Nasonia* oosome and surrounding cytoplasm are related to the ER, and thus likely to be RAVs as recently defined [35], we used an antibody against the KDEL peptide. This sequence is a highly conserved feature of ER resident proteins throughout animals that is commonly used to characterize ER shape and distribution. KDEL-labelled structures were punctate (Figure 5B-B’), consistent with the appearance of RAVs when observed with fluorescent microscopy [35]. Consistent with the vesicles observed by TEM (Figure 4A’’ and 5A), these punctate KDEL positive structures were also widely distributed throughout the oosome and the surrounding cytoplasm (Figure 5B-B’). To determine if RAVs were enriched in the oosome relative to the general cytoplasm we compared RAV count in segmented regions within and just outside of the oosome. We found no significant difference in RAV number between the oosome and the surrounding cytoplasm (Figure 5C).

**Figure 5:**
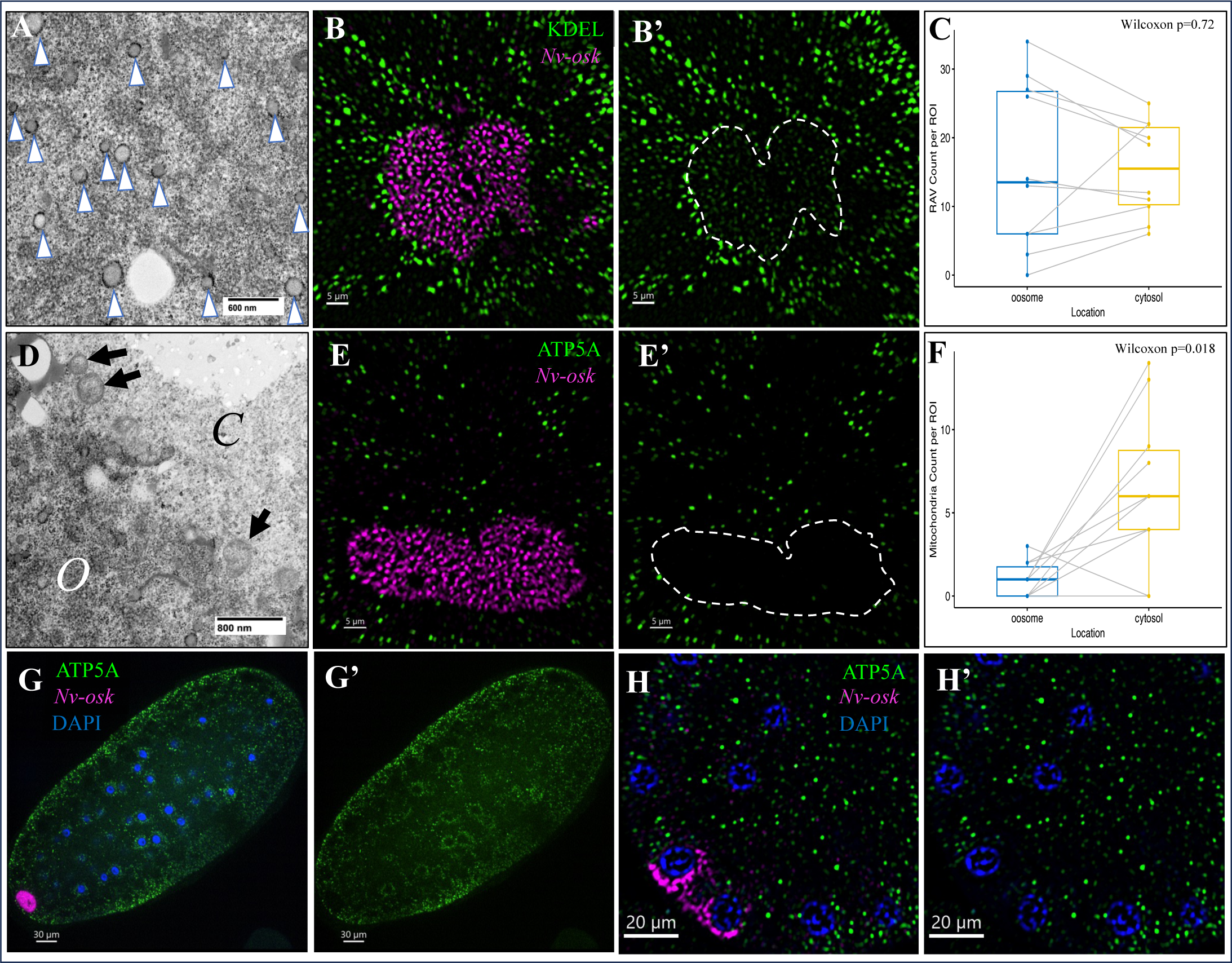
Oosome organelle interactions. **(A)** Electron micrograph of the oosome interior. Widespread RAV distribution marked with white arrowheads. Scale bar 600nm. **(B-B’)** Single plane confocal SRRF image showing the distribution of RAVs within and surrounding the oosome. *Nv-osk* (magenta) marks the oosome and RAVs (green) are labeled with KDEL. Scale bars 5 μm. **(C)** Boxplots of RAV count within a segmented region of equal size both inside and just outside of the oosome. Lines between indicate pairs from the same embryo. No significant difference determined by the Wilcoxon signed rank test. **(D)** Electron micrograph of the oosome periphery. The letter O indicates the oosome and C indicates the cytosol. Black arrows mark the mitochondria at the oosome periphery. Scale bar 800nm. **(E-E’)** Single plane confocal SRRF image showing the distribution of mitochondria within and surrounding the oosome. *Nv-osk* (magenta) marks the oosome and mitochondria (green) are labeled with anti-ATP5A. Scale bars 5 μm. **(F)** Boxplots of mitochondria count within a segmented region of equal size both inside and just outside of the oosome. Lines between indicate pairs from the same embryo. Significant difference determined by the Wilcoxon signed rank test. **(G-G’)** Single plane confocal image showing the enrichment of mitochondria (green) around nuclei (blue). Scale bar 30 μm. **(H-H’)** Single plane confocal SRRF image of decreased mitochondria (green) surrounding nuclei (blue) incorporated into the ooosme (magenta). Scale bar 20 μm. For both organelle counts n=10 embryos.

Unlike RAVs, while mitochondria were also apparent in the surrounding cytoplasm and at the oosome periphery, they appeared to be excluded from the oosome (Figures 4A’’&5D). To further validate this observation, we used the marker ATP5A to fluorescently label mitochondria and again compared the count in segmented regions both within and just outside of the oosome (Figure 5E-F). Consistent with our TEM observations, there are significantly fewer mitochondria inside the oosome than in the surrounding cytoplasm (Figure 5F). This finding is unexpected as germ plasm is generally enriched in mitochondria [30, 40, 41]. In *Drosophila*, the long isoform of Oskar, which is absent in *Nasonia* [8], is responsible for anchoring mitochondria at the posterior cortex [19]. The observed lack of mitochondrial enrichment suggests there is no equivalent mechanism for concentrating mitochondria in the region of pole cell formation in *Nasonia.* In fact, we suspect that the opposite may be the case. At later stages, mitochondria are highly concentrated around the pre-blastoderm syncytial nuclei (Figure 5G). When the presumptive pole cell nuclei enter the oosome at the pole cell budding stage, the vast majority of the mitochondria are removed from the nuclear periphery (Figure 5H’ and Movie S1). Based on these observations, the oosome may play a role in establishing a bottleneck of mitochondria at the very beginning of germline establishment and may thus be important in maintaining mitochondrial quality over generations.

Taken together, these findings show that while the oosome does contact and incorporate organelles into its internal network structure, there is some unknown means of discrimination whereby mitochondria are excluded. Further experiments are necessary to understand how mitochondria are excluded, and whether RAV incorporation is a passive part of oosome assembly or if it may play a functional or structural role.

## Discussion

Our analysis of the structure and composition of the oosome revealed numerous novel and unexpected features that highlight the striking diversity in membraneless organelle form. By investigating this exceptionally large and active germ plasm complex found in *Nasonia*, we broaden our understanding of which structural features may contribute to the stability of large condensates and of germ plasm evolution in insects.

### mRNA clustering in the oosome

Over the last several years, self-sorting of RNAs into homotypic clusters within germ granules has emerged as a surprisingly well conserved feature of germ granules across animal phyla [16, 17, 28, 29]. Our results partially conform to this trend, and potentially extend the forms homotypic RNA interactions can take. The scattered punctate distribution and lack of co-localization of low and moderately enriched mRNAs like *Nv-innexin* and *Nv-qin* is strongly suggestive of homotypic clustering of mRNA within the oosome environment (Figure 2B’, C’, &E). In *Drosophila*, mRNAs accumulate into homotypic clusters only in the context of core germ granules (composed largely of Osk and Vas protein) [27]. In contrast, the *Nv-qin* and *Nv-innexin* puncta do not appear to form in the context of a Nv-Osk and Nv-Vas complex. Rather they are scattered along an apparent network of highly enriched mRNAs of *Nv-bark* and *Nv-osk*.

The network distribution of *Nv-osk* and *Nv-bark* does not preclude homotypic interaction of these mRNAs. Indeed, the degree of colocalization between *Nv-bark* and *Nv-osk* is quite high (PCC=0.71), but somewhat lower than the value for the colocalization for two *Nv-osk* probe sets (PCC=0.87) (Figure 2D). This suggests that any overlap may be partial, and we cannot exclude that the apparent colocalization is a result of extremely high concentration of these molecules in a constrained area. To distinguish whether the *Nv-bark* and *Nv-osk* mRNA networks are truly largely overlapping, or whether they are actually distinct and potentially homotypic networks, methods that can further enhance the ability to resolve these tightly packed molecules, such as expansion fluorescence *in situ* hybridization (ExFISH), will be required [42].

### Potential origin and functional significance of the mesh-like mRNA network

Unlike the discrete puncta of moderately enriched mRNAs, the large-scale, mesh-like network distribution of *Nv-osk* and *Nv-bark* is not a pattern previously observed for embryonic germ granule mRNA. However, it was recently found that during late oogenesis in *Drosophila*, *oskar* particles “stick” together forming larger agglomerates [43]. While these are not on the scale of the oosome network, it does indicate the propensity of *oskar* particles to adhere to each other. This suggests the presence of a mechanism in *Nasonia*, potentially involving its close association with *Nv-bark* mRNA, further stabilizes these interactions allowing the formation of this large network. Additionally, a recent large scale RNAi screen in *Drosophila* identified nine endoplasmic reticulum (ER), nuclear pore complex (NPC), and cytoskeleton related proteins whose knockdown resulted in branched mesh-like morphology of polar granules, demonstrating the capacity of these granules to also form similarly shaped networks [20]. Considering the shared core assembly proteins between polar granules and the oosome, further investigation into how the oosome interacts with the ER, NPC, and cytoskeleton and how this compares with polar granules may provide insight into the evolution of its novel form.

In *Drosophila*, *osk* mRNA is toxic to pole cells and undergoes targeted degradation prior to and during pole cell formation [24]. The specificity of *osk* degradation appears to arise from its segregation into separate germ granules called founder granules, which associate with decapping and degradation factors [24]. In *Nasonia,* it has been shown that *Nv-osk* is similarly rapidly cleared from pole cells [8]. and we expand on this finding to show that *Nv-osk* mRNA levels begin to decrease significantly during embryogenesis shortly before pole cell budding (Figure S1). This finding further supports that removing *osk* ortholog mRNA is a conserved requirement for germ cell development in insects. However, our results revealed no indication that there is a clear equivalent to the founder granules that specifically segregate *Nv-osk* mRNA, which is distributed at high levels in a network throughout the oosome. This network does not appear to be correlated with early mRNA degradation, as we found a high degree of co-localization with at least one other mRNA, *Nv-bark* (Figure 2D) which, unlike *Nv-osk*, is maintained at high levels beyond pole cells through to the late embryonic gonads [3]. Thus, there is likely to be a distinct mechanism for regulating *Nv-osk* mRNA.

The degradation of *osk* precedes the destabilization and disassembly of founder granules in pole cells [24]. The conserved timing of significant *Nv-osk* degradation shortly before and during pole cell formation (Figure S1) [8], suggests a potentially similar mechanism for the oosome disassembly that occurs during pole cell formation to allow germ plasm partitioning. Future work to determine if the *Nv-osk* RNA network plays a structural role in the oosome will require more sophisticated techniques than currently available in *Nasonia*, as we cannot currently isolate early roles in nucleating the oosome during oogenesis from a later role in stabilizing the large granule. Nor can we clearly disentangle a structural role of the mRNAs from those of their presumed protein products since RNAi will affect both.

### Oosome-organelle interactions

Germ granules are known to associate with membrane-bound organelles [33, 34]. In fact, Balbiani bodies, another germline RNP complex found in immature oocytes and comparable in size to the oosome (10-100 μm), encompass many membrane-bound organelles such as the endoplasmic reticulum, Golgi and especially mitochondria. Therefore, we were unsurprised in our TEM imaging of the oosome to find that it does contain many membrane-bound organelles (Figures 4 and 5A-C). Conversely, the striking lack of mitochondria was unexpected (Figure 5D-F). Mitochondria are generally enriched in germ plasm to facilitate maternal inheritance, a process which is mediated by the long isoform of Oskar in *Drosophila* [19]. Since this long Oskar isoform is absent from *Nasonia* [8], we initially suspected an alternative enrichment mechanism exists in the wasp’s oosome, or posterior polar region. However, while mitochondria seem to be distributed around the cortex of the embryo and around the syncytial nuclei (Figure 5G), they seem to be nearly completely excluded from the oosome, even as it begins incorporating nuclei (Figure 5H & Movie S1). Given that other organelles, even larger ones like nuclei and lipid droplets, seem to be easily incorporated into the oosome (Figure 4), we propose that the oosome is selectively permeable to organelles and some unknown mechanism exists within the oosome that specifically prevents mitochondrial incorporation. Further imaging of how mitochondria interact with the oosome both during assembly, budding, and pole cell formation would provide a more comprehensive view of this mechanism.

The inclusion of ribosome-associated vesicles (RAVs) in the oosome was also unexpected since they had not yet been described in insect embryos, although they are suspected to be widespread across different tissue and cell types [35]. A recent study found that RAVs promote local translation in secretory cells by transporting active bound ribosomes [38]. Active translation has been observed previously in germ granules [44, 45], but it is still unknown whether translation occurs within the oosome specifically, and whether RAVs are involved in translational regulation. These questions will be addressed by future studies investigating post-transcriptional regulation within the oosome.

### Material properties of the oosome

How the unique compositional features of the oosome are reflected in its material properties is difficult to directly probe without the establishment of live imaging in *Nasonia*. We did, however, gain some insight from identifying patterns which can be attributed to specific oosome behaviors in response to the external forces that drive its migration. The most irregularly shaped oosomes are found immediately after egg lay when it is initially still attached to the cortex, they become more spherical during migration, and finally flatten upon re-establishing contact with the posterior cortex (Figure 1B & C). These patterns suggest that the oosome is at least partially liquid-like as it deforms and does not hold a specific shape under pressure, relaxes into a more spherical shape upon release from this pressure, and wets the posterior cortex upon reassociation (Figure 1). Furthermore, just before budding, the oosome coats incoming nuclei displaying additional wetting behavior and a potential model for the incorporation of other oosome-associated organelles (Movie S1). Additionally, the diminishing satellite particle count supports a more transient liquid-like particle regardless of whether it occurs by dissolution or fusion (Figure 3C), both of which have been previously observed in a germ granule context in the liquid-like P granules in *C. elegans* [31]. Although these observations suggest that the oosome is more liquid-like, we cannot currently dismiss the possibility that the oosome may be hydrogel-like or a combination of the two. Condensate material properties are often more complex than a single feature and can even be multiphasic as is the case for P granules which have a liquid-like PGL protein core [31], stabilized by surrounding gel-like MEG-3 clusters [46].

## Conclusion

This work has identified several unique and unexpected features of the oosome. Some provide plausible explanations for how the oosome maintains its unusually large size and complexity, for example the branching network of *Nv-osk* and *Nv-bark* mRNA running throughout the oosome. Others give potential insights into the functional significance of the oosome, such as the presence of RAVs. Further research along these lines will provide valuable insights into the assembly and function of membraneless organelles in general. In addition, the complexity and investment of a very high concentration of biopolymers localized to the oosome raises the question of what the evolutionary significance of the oosome structure is. It seems to be a conserved feature of the superfamily Chalcidoidea to which *Nasonia* belongs, as it has been described from several genera spanning the most distantly related lineages [13, 47, 48]. Comparing the composition and structure of oosomes within this group could provide unique insights into the evolutionary aspects of membraneless organelles at different time scales.

## Methods

### Embryo collection and fixation for in situ hybridization and immunofluorescence

Adult females were allowed to lay eggs on *Sarcophaga bullata* pupae at 25 °C for 3.5 hours in a modified petri dish apparatus called a Waspinator [49]. Host puparia were then cracked and *Nasonia* embryos were transferred directly to a fixation solution (5 mL heptane, 2 mL 10% methanol-free formaldehyde) and fixed overnight at room temperature. Apart from embryos labeled with Nv-Tud, which were fixed for 2 hours at room temperature and then overnight at 4°C. After fixation the vitelline membrane was removed by affixing the membranes to double-sided tape, adding phosphate-buffered saline with Tween (PBST), and then gently pushing with the flat edge of a 27-gauge needle to release the embryo from the attached membrane as described previously [50]. Fixed, devitellinized embryos were kept in PBST at 4°C for less than a week until fluorescence *in situ* hybridization and/or immunofluorescence was performed.

### Single-molecule FISH (smFISH)

Custom Stellaris RNA FISH oligonucleotide probe sets labelled with CALFluor590 or Quasar670 were generated complementary to *Nv-oskar*, *Nv-innexin*, *Nv-qin* and *Nv-barkbeetle*. Stellaris FISH was performed on devitellinized embryos following the manufacturers protocol for *Drosophila* embryos (LGC Biosearch Technologies). Stellaris RNA FISH was carried out for Figures 2 and 3 in tandem with immunofluorescence (IF), where IF was performed first and followed by Stellaris FISH. After staining, embryos were mounted under #1.5 glass coverslips in Fluoromount-G Mounting Medium with DAPI from Southern Biotech.

Hybridization chain reaction (HCR) was utilized to label *Nv-osk* RNA for the preparation of the oosome z-stacks used for sphericity and satellite particle measurements in Figures 1 and 3, as well as to provide an oosome marker during the organelle staining in Figure 5. We followed the Molecular Instruments dHCR v3.0 protocol for whole-mount fruit fly embryos, with the *Nasonia*-specific modifications outlined by Taylor & Dearden, 2021 [51]. In images acquired for Figure 5 dHCR was performed first and followed by immunofluorescence against KDEL or ATP5A. All embryos were mounted under #1.5 glass coverslips in Fluoromount-G Mounting Medium with DAPI from Southern Biotech.

### Immunofluorescence

Immunofluorescence was carried out in combination with smFISH, either before Stellaris, or after dHCR (see above). Devitellinized embryos were first washed 3 x 5 min with PBS/0.1%TritonX-100 followed by 1 hour blocking in PBS.0.1%TritonX-100/10%Western blocking reagent. Primary antibody incubation was at 4 °C overnight. Primary antibodies against core oosome proteins (Nv-Osk, Nv-Tud, and Nv-Vas) are described in Kharel et al., 2024 [18], Antibodies against mitochondria (anti-ATP5A antibody [15H4C4]) and endoplasmic reticulum (anti-KDEL antibody [EPR12668]) were obtained from Abcam. Primary antibody concentrations were: 1:250 guinea pig anti-Nv-Osk, 1:500 rabbit anti-Tud, 1:250 rabbit anti-Vas, 1:200 rabbit anti-KDEL, and 1:500 mouse anti-ATP5A. Embryos were then washed 3 times for 20 minutes in PBS/0.1%TritonX-100, and then were blocked for 20 minutes in PBS/0.1%TritonX-100/10% Western blocking reagent. Then incubation with Alexa Fluor 488-labeled secondary antibodies (1:500) for 2 hours at room temperature. Finally, embryos were washed five times for 20 minutes in PBS/0.1%TritonX-100. Stained embryos were either mounted under #1.5 glass coverslips in Fluoromount-G Mounting Medium with DAPI from Southern Biotech or underwent additional staining with Stellaris FISH (see above).

### Confocal microscopy and image processing

Confocal imaging was performed using an Andor Revolution WD spinning disc microscope with an Olympus inverted stand and 60x 1.4 NA silicon oil immersion objective. Z-stacks for sphericity, satellite particle, and *oskar* mRNA intensity measurements were acquired with an iXon 888 EMCCD camera in Andor Fusion software at a scan size of 50 μm and an auto-step size of 0.22 μm. All other oosome imaging were acquired with an additional 1.6x magnifier lens in the optical path. Super-resolution images were obtained using the SRRF-Stream+ version 1.16.1.0 plugin to the Andor Fusion software. Super-resolution radial fluctuations (SRRF) is an algorithm which analyzes subpixel radial and temporal fluorescence intensity fluctuations across a large number of frames acquired at the same position, to produce a single image with 2 to 6-fold increase in resolution obtainable by the 1MP iXon camera [52, 53]. All SRRF images in this paper were acquired using 100 frames per image. Pixel shift correction of multi-channel SRRF images was performed using 100 nm TetraSpeck microspheres as alignment markers.

### TEM sample preparation and microscopy

TEM embryos were collected from 0-1 hour egg lays, and initially fixed as described for FISH. They were then devitellinized in PBS instead of PBST. They were then post-fixed in 2%PFA/2.5%GA/0.1M Sodium Cacodylate pH 7.4 buffer for two hours at room temperature followed by overnight at 4°C. These were then washed in sodium cacodylate buffer (pH 7.2-7.4) and post-fixed in buffered 1% osmium tetroxide for 2 hours. After several buffer washes, samples were dehydrated in an ascending concentration of ethanol leading to 100% absolute ethanol, followed by 2 changes in propylene oxide (PO) transition fluid. Specimens were infiltrated overnight in a 1:1 mixture of PO and LX-112 epoxy resin, and 2 hours in 100% pure LX-112 resin, and then placed in a 60 C degree oven to polymerize (3 days). For better results, specimens were embedded between 2 sheets of Aclar plastic (vs. traditional embedding molds), regions of interest were etched out with a razor blade and glue-mounted onto blank blocks for subsequent microtomy.

Semi-thin sections (0.5-1.0 μm) were cut and stained with 1% Toluidine blue-O to confirm the areas of interest (via LM). Ultra-thin sections (70-80 nm) were cut using a Leica Ultracut UCT model ultramicrotome, collected onto 200-mesh copper grids and contrasted with 3% uranyl acetate and Reynolds lead citrate stains, respectively. Specimen were examined using a JEOL JEM-1400 Flash transmission electron microscope (operating at 80 kV). Micrographs were acquired using an AMT Side-Mount Nano Sprint Model 1200S-B (and/or Biosprint 12M-B cameras), loaded with AMT Imaging software V.7.0.1.

### Quantification of sphericity

Oosome sphericity was measured for surfaces generated in Imaris around z-projections of *Nv-osk* mRNA, which was fluorescently labeled with HCR FISH as described above. Oosome projections were sorted into syncytial nuclear cycles 0-7 by counting nuclei stained with DAPI, and 20 oosomes per cycle were analyzed to produce the sphericity boxplot with significance relative to the earliest timepoint NC0 indicated. Significance was determined with a Kruskal-Wallis test followed by a pairwise comparison with Dunn’s test. From the same pool of surfaces, 10 oosomes from each nuclear cycle were randomly selected to display as a gallery plot generated in Imaris with sphericity values depicted by color warmth.

### Satellite particle counts

Satellite particle surfaces were generated in Imaris using the same *Nv-osk* mRNA z-projections described for sphericity measurements. A standardized ROI of the embryo posterior surrounding the oosome was captured with consistent acquisition settings. Satellite particle counts and sub-oosome number were measured using the surfaces generated in Imaris. Sub-oosome presence was identified manually and confirmed by comparing surface volumes to identify satellite particles ≥ 20% of their corresponding osome surfaces volume. Boxplots were generated from satellite particle count data for each nuclear cycle, and significance relative to the earliest timepoint NC0 indicated. Significance was determined with a Kruskal-Wallis test followed by a pairwise comparison with Dunn’s test. For each nuclear cycle, 20 embryos were analyzed for both satellite particle count and sub-oosome presence.

### Organelle count per ROI

Using Imaris software, regions of interest of 256x256 pixels were generated both within and just outside of 10 oosomes, with dHCR labelled *Nv-osk* mRNA as an oosome marker. Imaris spot detection was run within these segmented ROIs to label either punctate KDEL or punctate ATP5A signal. A pairwise Wilcoxon rank sum test was then carried out comparing spot detection within and just outside each oosome. Control images of embryos stained using only secondary antibodies and no primary antibodies detected no spots using the same acquisition and spot detection settings.

### Pearson’s correlation coefficient (PCC) analysis

PCC were determined using the Bioimaging and Optics Platform (BIOP) version of the JACoP Plugin in Fiji [54]. An ROI was manually generated around each oosome in a SRRF, pixel-shift corrected image. Automatic thresholding options were applied to sample images and Otsu thresholding was identified as most accurately reflecting the signal across all mRNAs and proteins and therefore applied to each image during co-localization analysis. PCC outputs were averaged across oosomes for each condition and used to generate bar graphs in Figure 2. Costes randomization [55] was applied to each image to evaluate the significance of co-localization. The number of oosomes analyzed for each pair were as follows: five for Osk/*innexin* and Vasa/*qin*; six for *osk/osk*, *osk/bark*, Osk/*qin*, and Vasa/*innexin*; eight for *innexin/innexin* and *innexin/qin;* eleven oosomes for *osk/qin*, *osk*/*innexin*, *osk*/Vasa, and *osk*/Osk; and twelve for *osk*/Tud.

## Acknowledgements

We thank Figen Seiler and the UIC RRC Electron Microscopy Core for help with transmission electron microscopy. This work was supported by the NIH National Institute of General Medical Sciences grant (R01GM129153) to J.A.L. and A.L.A.

