## Supplementary Figures for "Novel structure and composition of the unusually large germline determinant of the wasp Nasonia vitripennis"

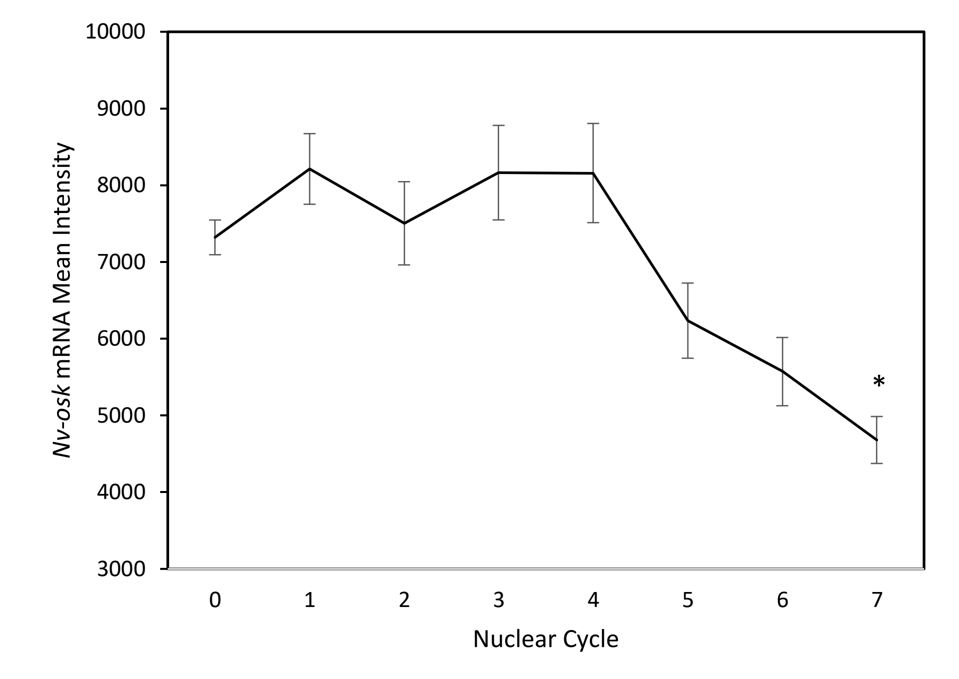


**Fig. S1 *Nv-osk* mRNA intensity decreases significantly prior to pole cell formation.** Values shown are mean ± S.E.M; n = 20 embryos per nuclear cycle Asterisks indicate statistical significance relative to before the first syncytial nuclear cleavage assessed by Dunn's Test.

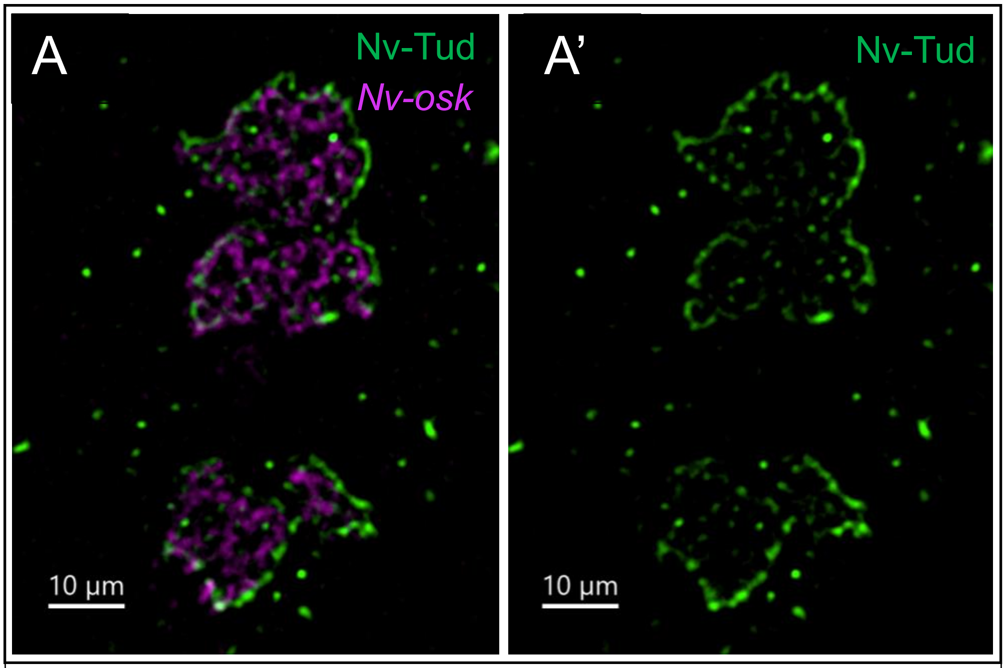


**Fig. S2 Large sub-oosomes can maintain the Tudor shell. (A)** A single frame of an oosome and sub-oosome with both the osk network (magenta) and Tudor (green) labeled. **(A')** A single frame an oosome and sub-oosome highlighting only the shared Tudor shell.
